# Complex interaction networks of cytokines after transarterial chemotherapy in patients with hepatocellular carcinoma

**DOI:** 10.1101/569939

**Authors:** Dong Wook Jekarl, Seungok Lee, Jung Hyun Kwon, Soon Woo Nam, Jeong Won Jang, Myungshin Kim, Yonggoo Kim

## Abstract

Inflammation in the tumor microenvironment influences all stages of HCC development and progression as well as the anti-cancer response by immune system. In this study, we studied cytokine networks before and after transarterial chemotherapy (TACE). Serum samples obtained from 203 HCC patients treated with TACE were analyzed for inflammatory cytokines including interleukin (IL)-1β, IL-2, IL-4, IL-5, IL-6, IL-9, IL-10, IL-12, IL-13, IL-17, IL-22, TNF-α, IFN-γ, and C-reactive protein (CRP) levels. Cytokine concentrations were measured at day 0 (D0, baseline), day3 (D3), day7 (D7), and day 60 (D60) after TACE. Network analysis revealed that modules within cytokine network at D0 were lost by D60 and modularity value (*Mc*) was decreased from 0.177 at D0 to −0.091 at D60. D60 had the lowest network heterogeneity and lower diameter, clustering coefficient, network density and recruited nodes. Degree correlation revealed that assortative network turned to disassortative network by D60 indicating that the network gained scale free feature. CRP, IL-2 were components of modules related with adverse outcome and IL-13, favorable outcome. Median survival month of patient group with high and low values with P-values were as follows: D0 CRP, 9.5 month (M), 54.2M (P<0.0001); D0 IL-2, 39.9M, 56.1M (P=0.0084); D3 CRP, 31.3M, 55.1 M (P=0.0056); D7 CRP, 28.7M, 50.7M (P=0.0065); IL-13, 51.9M, 33.6M (P=0.06). Network modularity decreased with temporal changes. Components of modules that included CRP, IL-2 and IL-6 were associated with adverse outcome and short overall survival. These modules were dissolved by D60 after TACE. Degree correlation decreased by D60, indicating that the cytokine network gained the scale free network property as in other biological network. TACE treatment converted cytokine network from that with inflammatory module to that with scale free network feature and without modules. Further studies are required to verify temporal changes of cytokine network in HCC patients after TACE.

## Introduction

Hepatocellular carcinoma is the most common malignancy of primary liver cancer, the 6th most commonly diagnosed cancer, and in 2018 was the fourth leading cause of cancer-related death worldwide [1]. In males, HCC is the 5th most common cancer and the 2nd leading cause of death, with an incidence rate twice that of females. Major risk factors for HCC are chronic hepatitis via hepatitis B and C viruses, alcoholic liver disease, aflatoxin B1-contaminated foods, nonalcoholic hepatitis, obesity, smoking, and type 2 diabetes [1, 2].

Pathophysiology of HCC is characterized by molecular gene alternations and driver mutations in the cell and the microenvironment surrounding the cell [3]; the inflammatory microenvironment caused by viral and toxic risk factors plays an important role in HCC development [4]. Chronically damaged tissues containing inflammatory molecules, fibrous tissue, and malignant cells construct a complex interaction network that promotes HCC development, progression, and resistance to therapy [3, 4]. The presence of nuclear factor-KB, epidermal growth factor, and interleukin (IL)-6 in the microenvironment is associated with poor prognosis [5]. Th2-like cytokines (including IL-4, IL-8, IL-10, and IL-5) are associated with aggressive and metastatic HCC, unlike Th1-like cytokines (including IL-1α, IL-1β, IL-2, TNF-α) [6]. TNF, lymphotoxin-α, lymphotoxin-β, and IL-6 are all associated with promotion of HCC [7, 8].

Because no single molecule acts as a lone factor for liver carcinogenesis, complex interaction networks should be considered. A network is comprised of a set of nodes that represent entities such as cytokines, genes, or proteins and a set of edges or links that define the relationships between nodes [9]. The relationship between nodes could be a physical interaction, physical link, ethereal connection, or represent mass/energy exchange [10]. A biological network can be composed of clustered molecules with both physical and non-physical interactions. Networks with physical interactions include protein to protein interaction networks, and networks with non-physical interactions include co-expression networks, disease networks, molecular pathway interactions, and gene versus phenotypes [11-12]. A conceptual network of cytokine co-expression data might reveal complex interactions in HCC patients treated with TACE. Therefore, in this study, a complex interaction network was explored by comparative analysis of multiple cytokines IL-1β, IL-2, IL-4, IL-5, IL-6, IL-9, IL-10, IL-12, IL-13, IL-17, IL-22, TNF-α, IFN-γ, and CRP, measured at baseline (D0), day 3 (D3), day 7 (D7), and day 60 (D60) after TACE.

## Materials and Methods

### Patients

This study was approved by the Institutional Review Board of Incheon St. Mary’s Hospital. After informed consent provided from participants, serum samples obtained from 203 HCC patients treated with TACE were analyzed for inflammatory cytokines including interleukin (IL)-1β, IL-2, IL-4, IL-5, IL-6, IL-9, IL-10, IL-12p70, IL-13, IL-17, IL-22, TNF-α, IFN-γ, and C-reactive protein (CRP) levels between June 2011 and December 2012 [13, 14]. For prognostic analysis and estimation of overall survival (six year), patient data were collected by December 2018. Patients with an unresectable tumor graded as Child-Pugh class A or B without evidence of portal vein involvement who were initially treated with TACE were selected [13]. HCC diagnosis was based on histology or elevated α-fetoprotein level in radiologic findings. TACE agents were composed of doxorubicin (50 mg) or epirubicin (50 mg) and cisplatin (60 mg) with lipiodol (5-10 mL) based on baseline tumor extent. Baseline characteristics of the datasets are listed in Table 1.

**Table 1.** Baseline patient characteristics.

| □ | Patient (n=203)□ |
| --- | --- |
| Age (year)□ | 58 (40 - 84)□ |
| Sex, Male / Female□ | 164 / 39□ |
| Etiology, HBV / HCV / Alcohol / Others□ | 142 / 17 / 22 / 22□ |
| Ascite, None / Present□ | 172 / 31□ |
| Portal hypertension, None / Present□ | 155 / 48□ |
| ECOG, 0 / 1 / 2 / 3 b□ | 115 / 49 / 27 / 4□ |
| Child Pugh Class, A / B / C b□ | 141 / 49 / 5□ |
| Tumor Maximum Size (cm)□ | 5.7 (0.8 - 28)□ |
| Tumor number, single / multiple□ | 102 / 101□ |
| Metastasis, None / present□ | 169 / 34□ |
| Treatment□ | □ |
| TAC / +PEI / +Radiation / sorafenib / Others□ | 127 / 3 / 16 / 5 / 52□ |
| Laboratory data, median (range)□ | □ |
| Platelet (x10<sup>9</sup>/L)□ | 131 (0.95 - 502)□ |
| Prothrombin time (INR)□ | 1.2 (1.0 - 461)□ |
| Bilirubin (mg/dL)□ | 1.2 (0.3 - 9.7)□ |
| AST (U/L)□ | 51.5 (11 - 522)□ |
| ALT (U/L)□ | 36 (11 - 392)□ |
| Albumin (d/dL)□ | 3.6 (0.8 - 5.1)□ |
| Creatinine (mg/dL)□ | 0.7 (0.4 - 1.3)□ |
| Sodium (mmol/L)□ | 138 (125 - 144)□ |
| AFP (ng/mL)□ | 57.6 (1.3 - 3,000)□ |
| PIVKA-II (mAU)□ | 193 (10 - 75000)□ |

This is the Table 1 legend.
HBV, hepatitis B virus; HCV, hepatitis C virus; TAC, transarterial chemotherapy; PEI, percutaneous ethanol infection; AST, aspartate aminotransferase; ALT, alanine aminotransferase; AFP, alpha fetoprotein; PIVKA-II, protein induced by vitamin K absence or antagonist II.
<sup>a</sup>The continuous variables are presented as median (range).
<sup>b</sup>For ECOG (Eastern Cooperative Oncology Group) and Child Pugh classification, 8 patient had no data.

### Cytokine measurement

The cytokines tested during TACE were IL-1β, IL-2, IL-4, IL-5, IL-6, IL-9, IL-10, IL-12, IL-13, IL-17, IL-22, TNF-α, and IFN-γ. Cytokines were measured using a Flowcytomix Multiplex with a multiple cytometric bead immunoassay (eBioscience, San Diego, CA, USA) at baseline (D0), day 3 (D3), day 7 (D7), and day 60 (D60) after TACE.

### Laboratory data measurements

CRP level, liver panel, and other blood chemistry data were measured at the same time as the cytokines using a Beckman Coulter AU5800 Clinical Chemistry System (Beckman Coulter, Miami, FL, USA). Data related to viral hepatitis were measured using anArchitecti2000 analyzer (Abbott Laboratories, Chicago, IL, USA). Platelet counts were measured using an XE-2100 differential analyzer (Sysmex Corporation, Kobe, Japan).

### Network topology analysis

Statistical analyses of cytokine profiles were performed with Spearman’s correlations, and statistically significant cytokine pairs were used for network analysis. Each cytokine was regarded as a node, and correlated pairs were regarded as links (edges) in the network [15]. Cytoscape and NetworkAnalyzer were used for network analyses [16-19]. Degree correlations, modularity and clustering analysis was calculated using iGraph which is a package of R program [20, 21].

As defined by graph theory, the clustering coefficient, <*C*>, is the number of edges from the nodes (cytokine), connected each other, which all allow for a connection to node *i*. The degree measurement is the number of edges (links or correlation pairs) connected to a node. Network density is the ratio of edges in the network to the total possible number of edges. The length of a path is the number of unique edges that form between two nodes. Distance is defined as the shortest path length between nodes *i* and *j* within the network. Network diameter is the maximum length of the shortest path. The characteristic path length is the average shortest path length that generates expected distance between two nodes, *i* and *j*. The average degree, <*k*>, indicates the average number of edges from the nodes. Network heterogeneity is a measure of variance in the number of edges divided by the mean number of edges. The topological coefficient is a relative measure for the extent to which a node shares a neighbor with another node. The network centralization value approaches 1 when the network resembles a star. Stress centrality of a node is the number of shortest paths passing through the node. Betweenness centrality of node *k* defines the shortest path between nodes *i* and *j* that pass through node *k* and implies that node *k* exerts control over other nodes. Closeness centrality of a node reflects how close it is to other nodes in the network. Eccentricity reflects maximum distance from a node [15-19]. Degree correlation (μ) is a value that node links to similar or dissimilar node, which was calculated by dividing degree correlation function divided by average number of degree. Modularity (*Mc*) was a fraction of links included within a given group compared to expected links randomly distributed which was calculated using random walks algorithm [22].

### Statistical analysis

Demographic data and baseline characteristics were presented as median values with ranges for continuous variables. The level of cytokines was compared among D0, D3, D7 and D60 using Kruskal-Wallis test. For molecules with statistical significance, Mann-Whitney test was performed. For the prognosis prediction, univariate and multivariate analyses were performed using Cox regression analysis, and all cytokine molecules were studied by entering the parameters into the model using backward method. Overall survival of patient outcome was analyzed by Kaplan-Meier method and compared using log rank method for parameters from multivariate model with statistical significance. The starting point of survival was the time of initial diagnosis of hepatocellular carcinoma and primary end point was death from any cause. The maximum area under the ROC curve (AUC) was selected for cut off values of parameters from multivariate analysis. Statistical analyses were performed using Medcalc software version 18.11 (Medcalc, Mariakerke, Belgium).

## Results

### Comparison of D0, D3, D7 and D60 cytokine levels and cytokine correlation analysis

The cytokine profiles on D0, D3, D7,and D60 after TACE showed that most of the cytokine concentrations were not statistically significant except CRP level, which was 19.45, 38.61, 32.62, and 26.94 mg/L, on D0, D3, D7, and D60, respectively, in Table 2 and S1 Fig. Except for the comparison between D0 and D60, all other comparisons among CRP levels were statistically significant by Mann-Whitney test. The IL-6 concentration increased over time from D0 to D60 and peaked at 56.16 ng/mL on D60, a 10-fold increase from the baseline concentration. Conversely, IL-2, IL-5, IL-10, IL-17α and IFN-γ concentration decreased over time and was the lowest on D60. IL-4, IL-13, TNF-α concentration was non-fluctuating or steady state on D60. IL-12, IL-6, IL-9, IL-22 and IL1-β increased on D60 after TACE.

**Table 2.** Measured cytokine levels for patients before transarterial chemotherapy (D0) and at 3 days (D3), 7 days (D7), and 60 days (D60) post-transarterial chemotherapy^a^.

|  | D0 |  | D3 |  | D7 |  | D60 |  |
| --- | --- | --- | --- | --- | --- | --- | --- | --- |
|  | Mean | SD | Mean | SD | Mean | SD | Mean | SD |
| IL-12 | 15.07 | 65.32 | 9.31 | 52.16 | 11.16 | 50.11 | 35.95 | 159.05 |
| IFN-γ | 6.19 | 39.28 | 4.74 | 26.24 | 8.13 | 34.61 | 13.51 | 50.79 |
| IL-17α | 19.97 | 78.14 | 12.2 | 40.19 | 11.88 | 41.57 | 20.43 | 101.35 |
| IL-2 | 26.75 | 50.29 | 30.03 | 43.22 | 26.5 | 49.46 | 31.5 | 75.96 |
| IL-10 | 2.79 | 9.86 | 3.35 | 13.48 | 2.82 | 9.95 | 6.78 | 29.27 |
| IL-9 | 5.99 | 27.44 | 4.81 | 25.57 | 6.25 | 26.13 | 19.71 | 77.41 |
| IL-22 | 156.46 | 278.54 | 98.59 | 176.21 | 128.24 | 238.72 | 268.58 | 518.17 |
| IL-6 | 4.14 | 14.3 | 15.05 | 46.61 | 10.18 | 28.14 | 12.13 | 31.67 |
| IL-13 | 32.97 | 33.53 | 30.75 | 36.48 | 31.02 | 33.24 | 42.17 | 43.93 |
| IL-4 | 22.39 | 53.86 | 21.6 | 50.34 | 22.3 | 51.39 | 41.21 | 95.03 |
| IL-5 | 2.83 | 21.16 | 1.85 | 13.48 | 0.92 | 6.47 | 0 | 0 |
| IL-1b | 13.22 | 49.69 | 11.27 | 39.72 | 12.27 | 34.12 | 29.4 | 95.7 |
| TNF-α | 11.5 | 34.63 | 13.72 | 50.1 | 7.72 | 22.32 | 19.95 | 85.72 |
| CRP<sup>a</sup> | 21.25 | 38.63 | 38.61 | 41.21 | 32.62 | 44.19 | 26.93 | 43.72 |

This is the Table 2 legend.
IL, interleukin; IFN, interferon; TNF, tumor necrosis factor; CRP, C-reactive protein; SD, standard deviation.
<sup>a</sup>Statistical analysis was performed using a Kruskal-Wallis test for analysis of variance among D0, D3, D7 and D60 and only CRP revealed statistical significance. Units for cytokines are as follows: CRP, mg/L; others, pg/mL.

Spearman’s method was performed to generate a co-expression matrix (S1-S4 Tables), and the number of molecular pairs significantly correlated at D0, D3, and D7, and D60 were as follows: D0, 44 pairs; D3, 58 pairs; D7, 56 pairs; and D60, 50 pairs. These data are listed in S5-S8 Tables.

### Network analysis of molecules

Networks were plotted for pairs of cytokines exhibiting significant correlations using correlation coefficients (Fig 1), and topological parameters were calculated (Table 3 and S9-S12 Table). D0, D3, D7, and D60 networks revealed that clustering coefficient, network density, and average number of neighbors were highest on D3. Network diameter, average shortest path and shortest path length were highest on D0 and equivocal or lowest on D60. Network heterogeneity and number of included nodes revealed decreasing patterns from D0 to D60, respectively. Modularity (*Mc*) of network revealed constantly decreasing pattern from D0 to D60 and the modules were colored in green or blue. Decreasing modularity values were also supported by clustering analysis of network (Fig 1). Degree correlation (μ) was converted from positive values to negative values by D60. At D0, μ was 0.281, at D3, μ was 0.209 and at D7, μ was 1.00. At D60, μ was −0.315, which implies that the network with assortative nature was turned to disassortative nature by D60 after TACE.

**Table 3.** Network topological parameters of D0, D3, D7, and D60 after transarterial chemotherapy.

|  | D0 | D3 | D7 | D60 |
| --- | --- | --- | --- | --- |
| Clustering Coefficient, $\langle C \rangle$ | 0.797 | 0.962 | 0.831 | 0.813 |
| Network density | 0.473 | 0.731 | 0.705 | 0.742 |
| Network heterogeneity | 0.398 | 0.338 | 0.378 | 0.307 |
| Connected components | 1 | 1 | 2 | 1 |
| Network diameter | 4 | 2 | 2 | 2 |
| Network centralization | 0.346 | 0.318 | 0.152 | 0.309 |
| Shortest path | 182 | 156 | 112 | 132 |
| Characteristic path length | 1.681 | 1.269 | 1.018 | 1.258 |
| Average degree, $\langle k \rangle$ | 6.143 | 8.769 | 8.462 | 8.167 |
| Number of nodes, $N$ | 14 | 13 | 13 | 12 |
| Degree correlation, $\mu$ | 0.281 | 0.209 | 1 | -0.315 |
| Modularity, $M_c$ | 0.177 | 0.038 | 0.036 | -0.091 |

**Fig 1.**
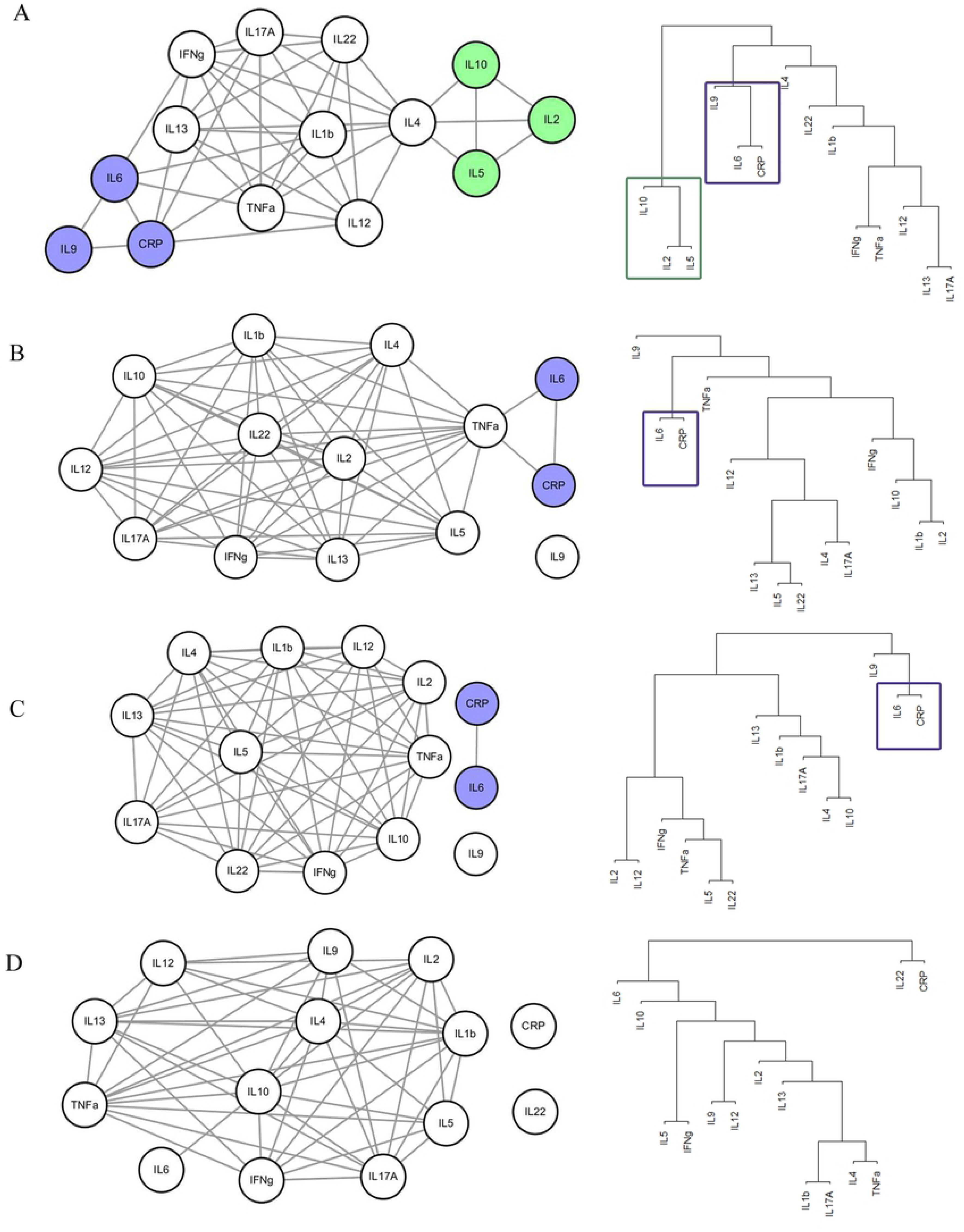
Cytokine Network of hepatocellular carcinoma after TACE. Network analysis results at (A) D0, (B) D3, (C) D7 and (D) D60 after transarterial chemotherapy are presented. Network modules are clustered and colored with blue or green.

Analysis of each node revealed that IL-4 had the highest degree (*k* = 10) on D0;TNF-α had the highest degree (*k* = 12) on D3;IL-1β, IL-2, IL-4, IL-5, IL-10, IL-13, IL-22, INF-γ and TNF-α had the highest degree on D7 (*k* = 10);and IL-10 had the highest degree (*k* = 11) on D60 (S9-S12 Tables). The node with the highest degree is considered the hub node or, in this study, a hub cytokine. IL-6 became weakly associated with other cytokines within the network and was linked with CRP on D7 andIL-10 on D60. IL-22 was strongly associated with network, which was isolated on D60. Alternatively, IL-9 was isolated from the rest of the cytokine network from D0, D3 and D7 on D60.

Altogether, network analysis revealed that smallest network diameter, decreased network heterogeneity and decreased modularity by D60 might be related to decreased cytokine network function. Loss of modularity was also supported by clustering analysis of network. Degree correlation showed that assortative network turned to disassortative network by D60 after TACE, which indicates that the network gained scale free feature.

### Prediction of patient outcome

For prediction of patient outcome, univariate and multivariate analysis was performed using all the studied molecules from D0, D3 (S13 Table) and D7, D60 (S14 Table). At D0, IL-2, IL-10, IL-6, IL-5, IL-1β, CRP were statistically significant in univariate analysis. In multivariate analysis, IL-2 revealed P-value of 0.012 with a hazard ratio (95% confidence interval, (CI)) of 1.003 (1.001-1.005). CRP revealed P-value of less than 0.001 with a hazard ratio (95% CI) of 1.017 (1.013-1.022). At D3, IL-17α, IL-6, IL-13, CRP revealed statistical significance and in multivariate analysis CRP revealed a P-value of 0.001 with a hazard ratio (95% CI) of 1.015 (1.006 - 1.024) and IL-6 revealed a P-value of 0.03 and a hazard ratio (95% CI) of 1.012 (1.001 - 1.022). At D7, only CRP revealed statistical significance in univariate analysis with a P-value of 0.016 with a hazard ratio of 1.009 (1.006-1.017). At D60, IL-12, IL-17α, IL-22, IL-6, IL-1β, TNF-α and CRP revealed statistical significance in univariate analysis. In multivariate analysis only CRP revealed statistical significance with a P-value of less than 0.001 with a hazard ratio (95% CI) of 1.022 (1.010 - 1.033). ROC analysis revealed that cut off values of D0 CRP was 8.87 mg/L and D0 IL-2 was 1.63 pg/mL. Cut off value of CRP at D3 was 22.39 pg/mL and IL-6 was 3.25 pg/mL. CRP at D7 was 16.27 mg/L and CRP at D60 was 7.53 pg/mL, respectively. Based on these parameters, overall survival was estimated. The five year survival of hepatocellular carcinoma with TACE patients was studied from the time of diagnosis to death of any cause. Mean survival month of patient group with high and low CRP concentration at D0 was 9.5 month (M) and 54.2 M, respectively, with a P-value of less than 0.001 (Fig 2). Of IL-2, patient group with high and low concentration at D0 revealed mean survival month of 39.9 M and 56.1 M, respectively, with a P-value of 0.0084. Mean survival month of CRP at D3 was 31.3M and 55.1 M, respectively, with a P-value of 0.0056. IL-6 at D3 revealed mean survival month of 33.7M and 53.1M for high and low concentration of IL-6 within patient group, respectively with a P-value of 0.0017. At D7, patient group with high and low concentration of CRP revealed 28.7 M and 50.7 M, respectively with a P-value of 0.0065.At D60, patient group with high and low concentration of CRP revealed mean survival month of 31.7M and 55.3M, respectively with a P-value of 0.0135.

**Fig 2.**
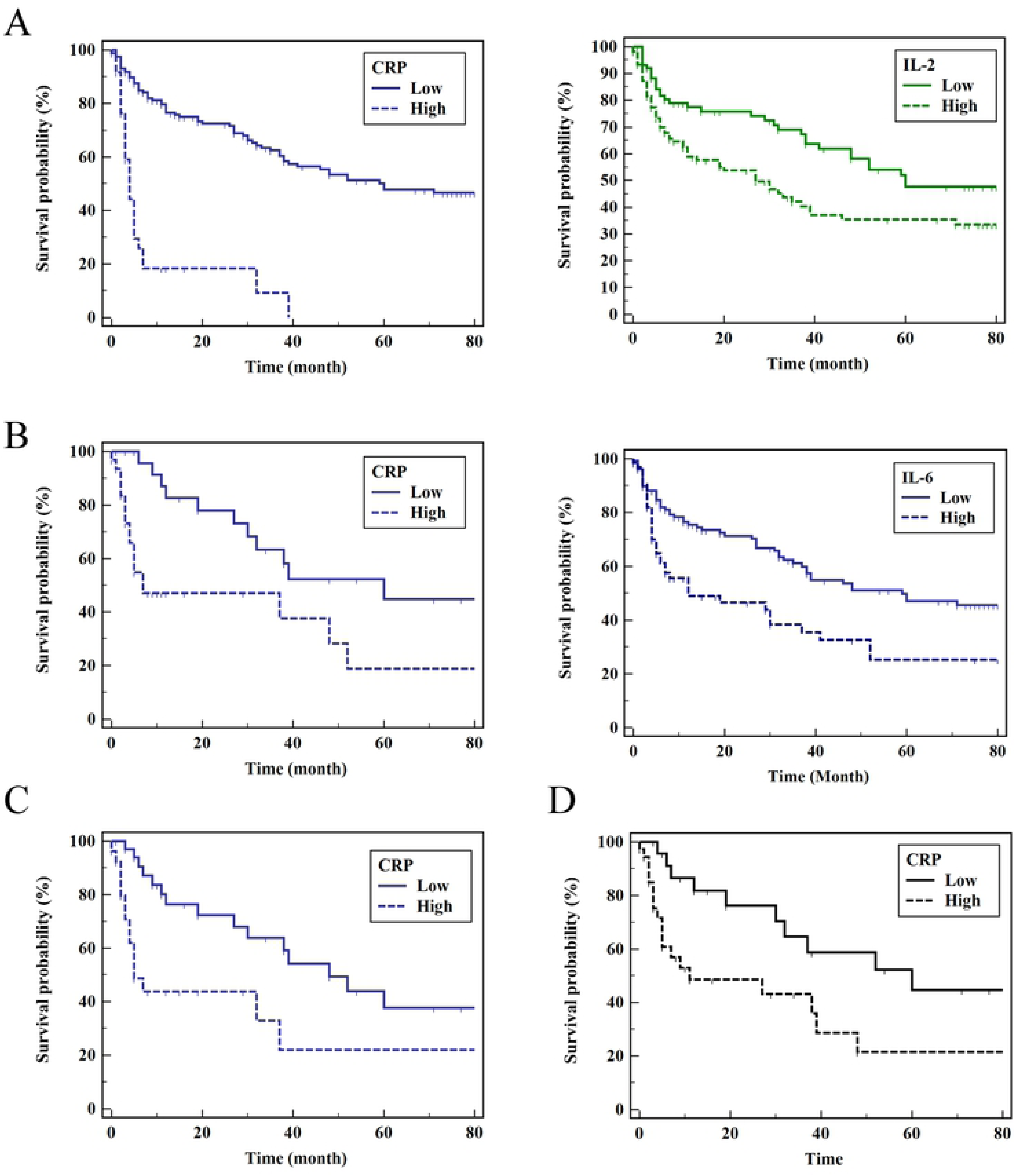
Patient outcome by cytokine status. Overall survival of patient group by high and low concentration of cytokines after transarterial chemotherapy was shown. (A) CRP and IL-2 at D0, (B) CRP and IL-6 at D3, (C) CRP at D7 and (D) CRP ant D60.

## Discussion

Carcinogenesis is characterized by genetic alteration of cells, and the tumor microenvironment is involved in all stages of carcinogenesis from initial cell transformation to metastasis [9]. The tumor microenvironment of HCC is composed of stromal cells, infiltrated innate and adaptive immune cells, and secreted cytokines in an extracellular matrix. An understanding of the complex network of interactions among these components could elucidate the steps of carcinogenesis and the influence of the microenvironment on hepatocytes. Cytokine molecules produced by innate and adaptive immune cells play an important role in the microenvironment of HCC. According to the Barcelona Clinic Liver Cancer (BCLC) staging system, patients with intermediate-stage HCC are generally treated with TACE, which reportedly lengthens median survival rates [23-25]. However, TACE often causes inflammation and ischemic injury to the liver and its effects on tumor microenvironment requires elucidation.

Understanding cytokine profiles may provide insight into immunological complexities in a tumor microenvironment influenced by treatment-associated inflammation, liver function, and HCC stage [13]. Comparison of cytokine concentrations among D0, D3, D7, and D60 were not statistically significant except for CRP, but most of the cytokine profiles revealed an increasing or decreasing pattern. IL-1β, IL-6, IL-9, IL-12p70, IL-22 level increased and IL-17α, IL-2, IL-5, IL-10, IL-17α and IFN-γ levels decreased, but IL-4, IL-13, and TNF-α remained stable by D60.

Cytokine profiles are complex, and network analysis might better elucidate different aspects of their dynamics that cannot be gleaned via single-cytokine analysis. The network heterogeneity, network diameter and network modularity helped explain the modular nature of the biological function [26]. Unlike for D0, the D60 cytokine network revealed decreased heterogeneity and diameter by D60 compared to D0, D3, and D7. These network parameters were supported by the modularity analysis that the modularity values constantly decreased from D0 to D60. At day D0 before TACE, there were 2 modules, which was colored in blue or green, and that was also shown by hierarchical clustering analysis of network (Fig 1). These parameters might imply that cytokine functions are decreased or normalized after TACE by D60.

Unexpectedly, survival analysis results were in line with network analysis. Multivariate analysis showed that CRP and IL-2 at D0, CRP, and IL-6 at D3, CRP at D7 and D60 was a predicting factor of adverse outcome. These molecules turned out to be one of components of modules in cytokine network at D0, D3 and D7 (Fig 1), except for D60 CRP. Therefore, the modules in cytokine networks are thought to be related with adverse outcome among patients and the modules bear unknown function that requires elucidation. In addition, response of TACE might be evaluated by reduction of tumor size and other measuring parameters as well as modularity (*Mc*) of the cytokine networks. Decreasing modularity from D0 to D60 implies that inflammatory aspects of cytokine network are decreased by TACE (Fig 2). At D0, component of modules colored in blue was IL-6, CRP, and IL-9 and modules colored in green was IL-2, IL-10 and IL-5 (Fig 1). Among them, CRP and IL-2 were related with adverse outcome in survival analysis. At D3, IL-6 and CRP constituted a module that was colored in blue and both of them were related with adverse outcome in survival analysis. At D7, CRP and IL-6 formed a module and CRP was related with adverse outcome. Network modularity results and survival analysis implies that module are related with adverse outcome. As modules were decreased from D0 to D60, TACE might have converted cytokine network from a network that have adverse outcome to that of relatively favorable outcome. Modules are thought to have inflammatory function that characterizes the network with that module.

Network diameter, shortest path, and path length are related to a small world property, which implies that short path length is associated with fast response against external stimuli and adaptation to environmental change [27]. In this study, these parameters were decreased in cytokine networks by D60, indicating that cytokine networks are relatively efficient in spreading perturbations within the network and in reacting to changes caused by external conditions. In addition, degree correlation values were also decreased by D60, which implies scale free feature of cytokine network by D60. Biological networks or scale-free networks tend to exhibit higher network heterogeneity [28, 29]. However, in this study, network heterogeneity parameter was decreased at D60 compared to other data points, which was inconsistent with other parameters and was unexpainable. Altogether, cytokine network are thought to be converted from network with inflammatory function to decreased function after TACE.

It is of interest to see the weak association of IL-6, isolation of IL-22 and incorporation of IL-9 into the cytokine network on D60. IL-6 was correlated with tumor size, number, and metastasis and is an unfavorable prognostic indicator in HCC [13, 30]. Weak association of IL-6 on D7 and D60 indicates that either the expression level of IL-6 was uncontrolled by the cytokine network or negative feedback mechanism of expression of IL-6 was uncontrolled. These might be explained by forming modules with other molecules. IL-6 formed module with CRP from D0 to D7 and related with adverse outcome in D3. In contrast, IL-9 was incorporated into the cytokine network and IL-9 was isolated on D3 and D7. IL-9 is known to be produced by various CD4 T-cells including T_H_2, T_H_9, T_H_17, and T_reg_ [31]. IL-9 affects hematopoietic cells, epithelial cells, mast cells, smooth muscle cells, and lymphocytes and acts as an immunomodulator [32]. Further studies are required, as little is known about the role of IL-9 in HCC patients. IL-22 level was increased and was isolated from the cytokine network on D60. IL-22 has been reported to have protective properties for hepatocyte [33] or promoted cancer cell growth and metastasis [30, 31]. Cytokine network implies that IL-6 and IL-22 might have lost negative feedback mechanism after TACE.

A node with the highest degree could be regarded as a hub node, and hub nodes tend to link or interact with nodes with fewer edges [19, 35]. The hub node is related to network robustness, as removal of hub nodes might lead to system failure [27]. In this study, the hubs were IL-4 on D0; TNF-α onD3; IL-1β, IL-2, IL-4, IL-5, IL-10, IL-13, IL-22, INF-γ, and TNF-α on D7; and IFN-γ, IL-2, and IL-10 was the hubs on D60.

The degree correlation is a parameter that defines that relation of nodes with other nodes. Assortativeness indicates that hub is related with hub and dissorativeness indicates the hub node is related with non-hub node, which is close to hub and spoke model. In this study, the positive value of degree correlation value (μ) changed to negative after chemotherapy by D60. TACE treatment changed assortative network to disassortative network by D60. Considering that most of biological networks possess disassortative feature, TACE treatment turned cytokine network into network similar to normal biological network.

Limitations of this study are that the etiology of HCC in this study was mostly due to hepatitis B virus, which might have a different microenvironment compared to other causes and the patients were predominantly male, which might have affected the outcome. In addition, the cytokine network might have been better analyzed using a time series algorithm that was mostly designed for analysis of gene expression data. Comparisons of network topological parameters using Z scores were statistically impossible due to the use of four networks.

## Conclusion

In conclusion, cytokine network of HCC patients before and after TACE revealed different features. Network modularity decreased with temporal changes. Components of modules that included CRP, IL-2 and IL-6 were associated with adverse outcome and short overall survival. These modules were dissolved by D60 after TACE. Degree correlation decreased by D60, indicating that the cytokine network gained the scale free network property as in other biological network. TACE treatment converted cytokine network from that with inflammatory module to that with scale free network feature and without modules. Further studies are required to verify these temporal changes of cytokine network in HCC patients after TACE treatment.

## Acknowledgements

We thank to all the participants in this study.

## Supporting information

**S1 Fig. Bar graph of cytokines for day 0, 3, 7 and 60**. Median and 95% confidence interval (CI) were presented.

**S1 Table. Correlation matrix of cytokines concentrations at D0.**

**S2 Table. Correlation matrix of cytokines concentrations at D3.**

**S3 Table. Correlation matrix of cytokines concentrations at D7.**

**S4 Table. Correlation matrix of cytokines concentrations at D60.**

**S5 Table. P-value of correlation matrix from cytokines concentrations at D0.**

**S6 Table. P-value of correlation matrix from cytokines concentrations at D3.**

**S7 Table. P-value of correlation matrix from cytokines concentrations at D7.**

**S8 Table. P-value of correlation matrix from cytokines concentrations at D60.**

**S9 Table. Topological parameters from network analysis of D0.**

**S10 Table. Topological parameters from network analysis of D3.**

**S11 Table. Topological parameters from network analysis of D7.**

**S12 Table. Topological parameters from network analysis of D60.**

**S13 Table. Univariate and multivariate analysis by Cox regression analysis for D0 and D3.**

**S14 Table. Univariate and multivariate analysis by Cox regression analysis for D7 and D60.**

## References

1. Bray F, Ferlay J, Soerjomataram I, Siegel R, Torre L. Global cancer statistics 2018: GLOBOCAN estimates of incidence and mortality worldwide for 36 cancers in 185 countries. CA Cancer 2018;68:394–424.

2. Sanyal AJ, Yoon SK, Lencioni R. The etiology of hepatocellular carcinoma and consequences for treatment. The Oncologist. 2010;15(Supple4):14–22.

3. Llovet JM, Zucman-Rossi J, Pikarsky E, Sangro B, Schwartz M, Sherman M, et al. Hepatocellular carcinoma. Nat Rev Dis Primers. 2016;2:1–23. doi:10.1038/nrdp.2016.18.

4. Hernandez-Gea V, Toffanin S, Freidman SL, Llovet JM. Role of the microenvironment in 2. 527.

5. Hoshida Y. Gene expression in fixed tissues and outcome in hepatocellular carcinoma. N Engl J Med. 2008;359:1995–2004.

6. Budhu A, Forgues M, Ye QH, Jia HL, He P Zanetti, et al. Prediction of venous metastases, recurrence, and prognosis in hepatocellular carcinoma based on a unique immune response signature of the liver microenvironment. Cancer Cell. 2006;10:99–111.

7. Bauer J, Namineni S, Reisinger F, Zoller J, Yuan D, HeikenWalder M. Lymphotoxin, NF-KB and cancer: the dark side of cytokines. Dig Dis. 2012;30:453–468.

8. Taniguchi K, Karin M. IL-6 and related cytokines as the critical lynchpins between inflammation and cancer. Semin Immunol. 2014;25:54–74.

9. Estrada E. Quantifying network heterogeneity. Phys Rev. 2010;82:0660102-1-066012-8.

10. Estrada E. Evolutionary equations with application in natural sciences. Introduction to complex network: structures and dynamics. Switzerland: Springer Nature; 2015.

11. Barabasi AL, Oltvai Z. Network biology: understanding the cell’s functional organization. Nat Rev Genet. 2004;5:101–113.

12. Barabasi AL, Gulbahce N, Loscalzo J. Network medicine: a network-based approach to human disease. Nat Rev Genet. 2011;12:56–68.

13. Kim MJ, Jang JW, Oh BS, Kwon JH, Chung KW, Jung HS, et al. Change in inflammatory cytokine profiles after transarterial chemotherapy in patients with hepatocellular carcinoma. Cytokine. 2013;64:516–522.

14. Jang JW, Oh BS, Kwon JH, You CR, Chung KW, Kay CS, et al. Serum interleukin-6 and C-reactive protein as a prognostic indicator in hepatocellular carcinoma. Cytokine. 2012;60:686–693.

15. Jekarl DW, Kim KS, Lee S, Kim M, Kim Y. Cytokine and molecular networks in sepsis cases: A network biology approach. Eur Cytokine Netw 2018;29:103–111.

16. Doncheva NT, Assenov Y, Domingues FS, Albrecht M. Topological analysis and interactive visualization of biological networks and protein structures. Nat Protoc. 2012;7:670– 685.

17. Saito R, Smoot ME, Ono K, Ruscheinski J, Wang P, Lotia S, et al. A travel guide to Cytoscape plugins. Nat Method. 2012;9:1069–1076.

18. Netanalyzer [Internet]. Munchen: Max planck Societ; [cited 15 APR 2018]. Available from http://med.bioinf.mpi-inf.mpg.de/netanalyzer/.

19. Zhu Z, Gerstein M, Snyder M. Getting connected: analysis and principles of biological networks. Genes Dev. 2007;21:1010–1024.

20. Barabasi AL. Network science. United Kingdom: Cambridge University Press; 2016.

21. Csardi G, Nepusz T. The igraph software package for complex network research, InterJournal, Complex Systems. 2006;1695–1702.

22. Newman M. Modularity and community structure in networks. Proc Natl Acad Sci. USA. 2006;103:8577–8582.

23. Llovet. JM. Briux J. Systematic review of randomized trials for unresectable hepatocellular carcinoma: chemoembolization improves survival. Hepatology. 2003;37:429–442.

24. European association for the study of the liver and European organization for research and treatment of cancer. EASL-EORTC clinical practice guidelines: management of hepatocellular carcinoma. J Hepatol. 2012;56:908–943.

25. Xue T, Jia Q, Ge N, Chen Y, Zhang B, Ye S. Imbalance in systemic inflammation and immune response following transarterial chemoembolization potential increases metastatic risk in huge hepatocellular carcinoma. Tumor Biol. 2015;36:8797–8803.

26. Pavlopoulos GA, Secrier M, Moschopoulos CN, Soldatos TG, Kossida S, Aerts J, et al. Using graph theory to analyze biological networks. BioData Mining. 2011;4:10–37.

27. Boccaletti S, Latora V, Moreno Y, Chavez M, Hwang DU. Complex networks: structure and dynamics. Phys Rep. 2006;424:175–308.

28. Jacob R, Harikrishnan KP, Misra R, Ambika G. Measure for degree heterogeneity in complex networks and its application to recurrence network analysis. R Soc open Sci. 2017;4:160757.

29. Cheng F, Liu C, Shen B, Zhao Z. Investigating cellular network heterogeneity and modularity in cancer: a network entropy and unbalanced motif approach. BMC Sys Biol. 2016;65(Suppl 3):302–311.

30. Porta C, De Amici Quaglini S, Paglino C, Tagliani F, Boncimino A, et al. Circulating interleukin-6 as a tumor marker for hepatocellular carcinoma. Ann Oncol. 2008;19:353–358.

31. Noelle RJ, Nowak EC. Cellular sources and immune functions of interleukin-9. Nat Rev Immunol. 2010;10:683–687.

32. Goswami R, Kaplan M. A brief history of IL-9. J Immunol. 2011;186:3283–3288.

33. Park O, Wang H, Weng H, Feigenbaum L, Li H, Yin S, et al. In vivo consequences of liver-specific interleukin-22 expression in mice: implications for human liver disease progression. Hepatology. 2011;42:252–261.

34. Jiang R, Tan Z, Deng L, Chen Y, Xia Y, Gao Y, et al. Interleukin-22 promotes human hepatocellular carcinoma by activation of STAT3. Hepatology. 2011;54:900–909.

35. Xu Ke, Bezakova I, Bunimovich L, Yi SV. Path lengths in protein-protein interaction networks and biological complexity. Proteomics. 2011;11:1857–1867.

